# Listening to speech induces coupling between auditory and motor cortices in an unexpectedly rate-restricted manner

**DOI:** 10.1101/148288

**Authors:** María Florencia Assaneo, David Poeppel

## Abstract

The relation between perception and action remains a fundamental question for neuroscience. In the context of speech, existing data suggest an interaction between auditory and speech-motor cortices, but the underlying mechanisms remain incompletely characterized. We fill a basic gap in our understanding of the sensorimotor processing of speech by examining the synchronization between auditory and speech-motor regions over different speech rates, a fundamental parameter delimiting successful perception. First, using MEG we measure synchronization between auditory and speech-motor regions while participants listen to syllables at various rates. We show, surprisingly, that auditory-motor synchrony is significant only over a restricted range and is enhanced at ~4.5 Hz, a value compatible with the mean syllable rate across languages. Second, neural modeling reveals that this modulated coupling plausibly emerges as a consequence of the underlying neural architecture. The findings suggest that the auditory-motor interaction should be interpreted rather conservatively when considering phase space.

## INTRODUCTION

The relation between perception and action has been investigated extensively and debated vigorously, in neuroscience as well as psychology. A particularly prominent example investigating this relation comes from research on speech. One line of study, refreshingly, has yielded converging evidence: when examining possible perception-production links from the perspective of speech *production*, there is emerging consensus that there exists a tight, mechanistic, causal link between the speech-motor output systems and the auditory-perceptual systems that underpin the monitoring and guidance of speech motor control. Note that a link to the somatosensory system, likewise, is well supported (Tremblay et al. 2003). This direction of interaction is theoretically well motivated and computationally explicit (Hickok et al. 2011; Guenther 2016), as well as attested in a variety of neural data (Cheung et al. 2016; Wilson et al. 2004; Tian & Poeppel 2012). The motor-to-auditory coupling, well summarized as the sensory guidance of speech-motor control, has been shown to provide plausible functional roles across different tasks. Examples include (i) establishing motor-acoustic mappings during learning and development (Kuhl 2010; Byun et al. 2013) and (ii) prediction of corresponding speech signals from motor plans (Rauschecker & Scott 2009; Tian & Poeppel 2012) thereby enabling online corrections of the speech motor-commands by comparing the predictions with the actual incoming auditory feedback (Behroozmand & Larson 2011; Houde & Nagarajan 2011; Guenther 2016).

On the other hand, considerable controversy arises concerning the role that the speech-motor system plays during perception. A body of evidence from neuroscience documents a link between perceptual and motor systems under diverse speech perception circumstances. For example, experiments show that (i) during the processing of speech (Wilson et al. 2004; Cheung et al. 2016), especially in the presence of noise (Du et al. 2014), speech-motor areas are active; (ii) transcranial magnetic stimulation of motor cortex interferes with phonological discrimination tasks (Meister et al. 2007; Sato et al. 2009; Möttönen et al. 2012); and (iii) induced mechanical modifications of vocal tract articulators influence speech perception in adults (Ito et al. 2009) and infants (Bruderer et al. 2015). It is noteworthy, however, that consensus has not been reached on which features of the speech sounds might be encoded in motor activation. Some experiments report a somatotopic activation of motor areas according to the place of articulation of the perceived speech sounds (Pulvermüller et al. 2006; Correia et al. 2015); others do not reproduce such patterns, showing rather different encoding schemes (Arsenault & Buchsbaum 2015; Cheung et al. 2016). Moreover, while some claims -in line with the motor theory of speech perception (Liberman et al. 1967)- argue that articulator-specific representations in motor areas are necessary when the corresponding input requires perceptual analysis (Pulvermüller & Fadiga 2010), others postulate the motor activation as an epiphenomenon that could at best assist perception in the case of degraded auditory signals (Lotto et al. 2009). Lesion evidence also calls into question the speech-motor system’s functional role (Bishop et al. 1990; Naeser et al. 1989). These controversies concerning the putative functional role of the motor system for speech perception may have masked questions about *how* an auditory to speech-motor cortex link might be achieved in the first place. The neurophysiological study we report takes on this issue from a new perspective to fill this basic gap. We explore how the link could be mechanistically achieved, regardless of a potential functional role. Specifically, we aim to understand a basic neurophysiological property that underpins how sensory and speech-motor regions interact. To test this, and to provide a new window onto a potential mechanism, we exploit the concept of synchrony (Buzsaki 2009) between auditory and speech-motor regions.

On the sensory side, it is well established that auditory cortex activity entrains to speech faithfully (Ahissar et al. 2001; Luo & Poeppel 2007) and that this brain-to-signal synchronization is necessary to produce auditory representations that yield intelligible speech (Lakatos et al. 2007; Henry & Obleser 2012; Peelle et al. 2013; Doelling et al. 2014). This neural entrainment is especially strong in the theta frequency band (4-7 Hz). Importantly, the time scale associated with theta-band activity aligns closely with standard speech envelope modulation rates, which in turn corresponds to cross-linguistically typical syllable rates (Ding et al. 2017; Pellegrino et al. 2011). Is the robust coupling between acoustic stimuli and auditory cortical activity linked to the sensorimotor machinery? Recent results using electrophysiological approaches provide valuable first steps in elucidating how synchronized activity can play a role in facilitating communication across regions during speech comprehension (Alho et al. 2014; Park et al. 2015). These experiments, although their focus lies on comprehension and not mechanism, demonstrate that inter-areal synchrony might support sensorimotor integration. However, the key properties have not been characterized, in particular the sensitivity to syllable rate, arguably the most fundamental property of speech perception and production.

Here we identify a critical aspect and basic architectural constraint of the auditory-motor circuitry for speech from the viewpoint of synchrony. Using magnetoencephalography (MEG) recordings, we measure how the coupling between auditory and motor regions, identified in individual participants’ brains, is modulated while they listen to syllables presented at various rates. The simplest hypothesis about how auditory and speech-motor regions interact is that they are coupled equally across all (plausible) speech rates. Surprisingly, we find that auditory-motor coupling, as quantified by inter-areal neural synchrony, is only significant over a restricted range and shows a specific enhancement at ~4.5 Hz, the scale that aligns closely with the mean syllable rate across languages. Computational analyses reveal that this restricted synchronization could emerge as a consequence of the underlying neural architecture. Numerical simulations in which motor cortex is represented by inhibitory and excitatory synaptically-coupled neural populations (Wilson & Cowan 1972) reproduce the experimental synchronization findings. Despite the long history of exploring the link between speech-motor and auditory areas, few experiments have tested in a principled manner how coupling may be mechanistically achieved. Our results contribute one new step along the way.

## RESULTS

We designed an MEG protocol to characterize sensorimotor integration in speech processing as a function of syllable rate. Participants were presented with a set of audio trials, consisting of trains of synthesized syllables played at different rates, while their brain activity was recorded. The signals originating in auditory and speech-motor cortices, derived from the raw magnetic field, were then submitted to further analyses.

Seventeen individuals participated; each completed two speech-based localizer conditions and then listened to the different auditory conditions of interest. Each stimulus consisted of 3 seconds of silence (baseline) followed by 6 seconds of syllable repetition. Syllables were repeated at five different rates: 2.5, 3.5, 4.5, 5.5 and 6.5 syllables per seconds (Figure 1A). To insure the participants’ attention to the stimuli, after each trial they indicated if a given target syllable was present.

**Figure 1.**
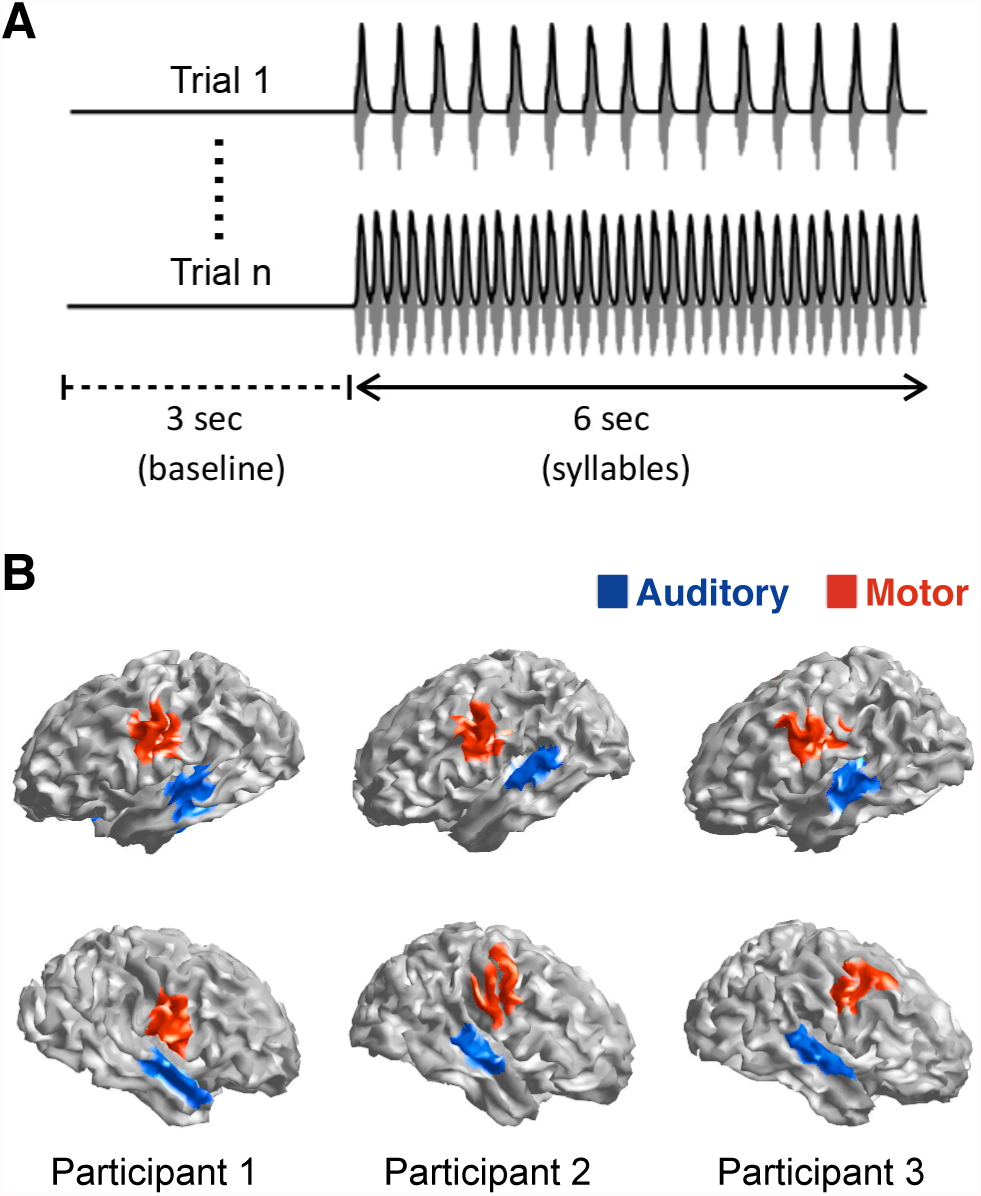
Extracting motor and auditory cortex activity while perceiving speech. A) Two examples of the experimental trials. Grey: sound wave. Black: its envelope. The upper trial shows a syllable rate of 2.5 Hz, the lower 6.5 Hz; trial order of the five conditions was randomized for each participant. B) Functional localizations of speech-motor (red) and auditory (blue) regions for three participants displayed to show the consistency and variability across individuals. Top row – left hemisphere; bottom row – right.

In order to extract the time series of the activity generated in specific cortical areas, participants first performed independent auditory and motor localizer tasks during the MEG session (see Methods). Additionally, each participant’s structural magnetic resonance image (MRI) was obtained, and we used source reconstruction techniques (MNE) to generate a reliable localization of bilateral auditory and speech-motor areas for each participant (Figure 1B). Finally, time series were obtained of the activity elicited in the main experiment (syllable presentation) in the localized areas of interest (see Methods).

### Activity in auditory cortices synchronizes to the speech envelope across rates

The experiment capitalizes on the well-established finding of entrainment of auditory cortical activity to the envelope of speech, resulting in an enhancement of brain-stimulus synchrony for frequencies around the perceived syllable rate (Ahissar et al. 2001; Doelling et al. 2014; Henry & Obleser 2012; Luo & Poeppel 2007). In order to confirm that the present data replicate this effect, and to explore if entrainment is modulated by syllable rate, we computed the phase locking values (PLV) between auditory cortex time series and the speech envelopes (see Methods).

A first characterization of the data reveals, as expected, an enhanced synchronization of the neural signal to the speech envelope at the frequency corresponding to a given syllable rate (Figure 2A). Each PLV plot shows a peak surrounding the rate of the stimulus (increment PLV relative to the 3- sec duration baseline). To quantify this result, we compute the mean PLV, per condition and participant, around the perceived syllable rate (Figure 2B; see Methods). First, for all five rate conditions, the PLV shows a significant increment from baseline (Wilcoxon signed rank test p<0.01, corrected for false discovery rate, FDR). Note that here we display the average across hemispheres; the PLV for individual regions (right and left auditory areas) did not contribute differentially (Figure S1). A Kruskal-Wallis test reveals a significant difference across conditions (χ^2^(4)=15.18, p<0.05). This effect appears to be driven by the weaker response profile at the 6.5 syllable rate condition. A post-hoc analysis comparing the central condition (4.5 syllables per second) against all others shows no statistical differences (Wilcoxon signed rank test, FDR-corrected, p_4.5|2.5_=0.25, p_4.5|3.5_=0.72, p_4.5|5.5_=0.72, p_4.5|6.5_=0.16).

**Figure 2.**
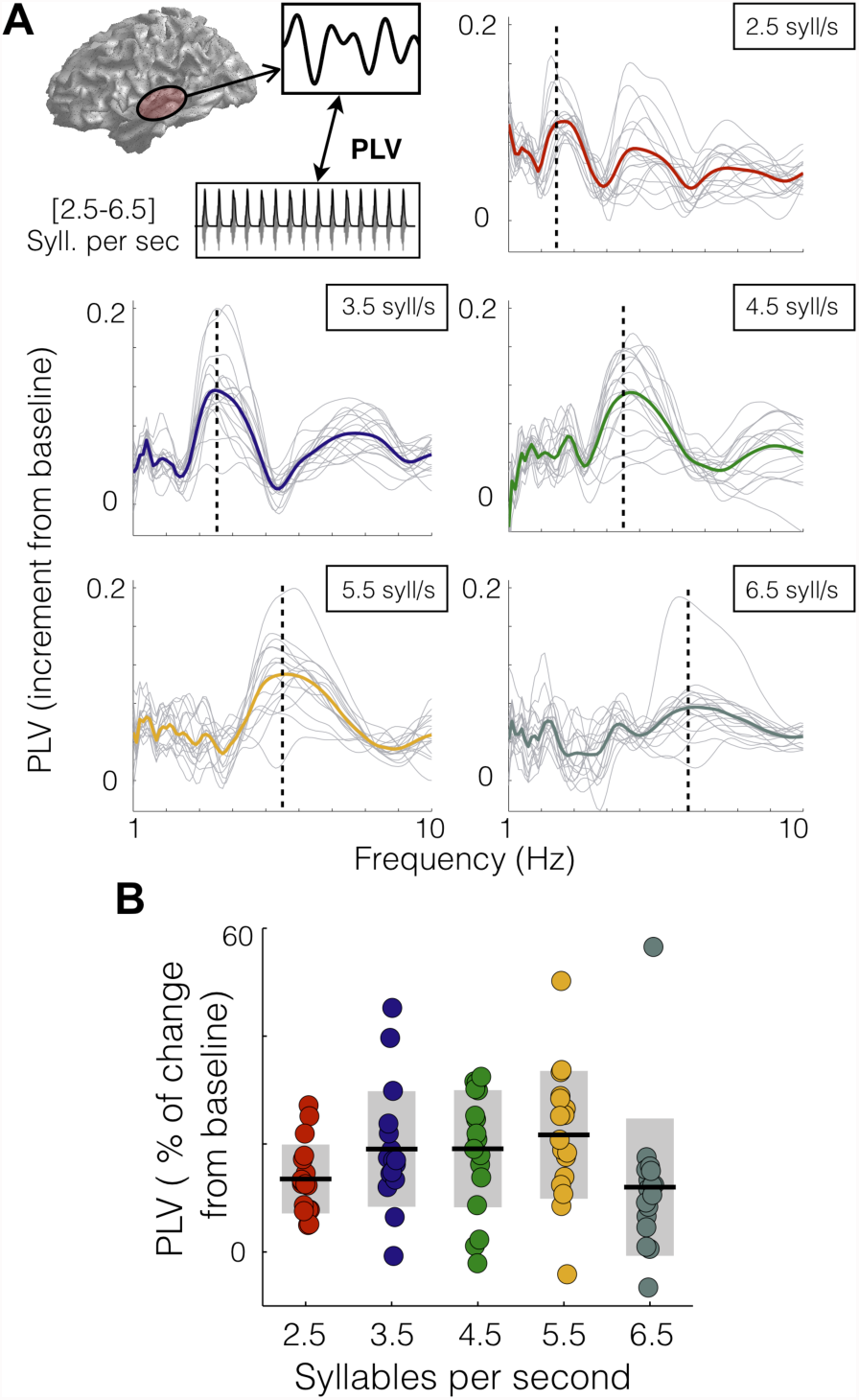
Activity in auditory cortices synchronizes to the speech envelope across rates. A) Increment from baseline of PLV between auditory cortex and the envelope of the sound as a function of frequency. Panels correspond to different syllable rate conditions. Light gray lines: individual subject data. Colored line: mean across subjects (red: 2.5, blue: 3.5, green: 4.5, yellow: 5.5 and dark gray: 6.5 syllables per seconds). Vertical line: stimulus rate. B) Mean PLV around the syllable rate of each condition (syll. rate +/- 0.5 Hz). Dots: individual participants. Black lines: mean across participants; shadowed region is standard deviation.

### Synchronization between auditory and motor cortices reveals a preferred rate

We next examined the sensorimotor integration of speech. A range of studies implicate the activation of motor cortex while listening to syllables (Du et al. 2014; Pulvermüller et al. 2006; Wilson et al. 2004), but it is worth recalling that this putative sensorimotor linkage remains controversial. Here we pursue a new perspective, testing the existence of phase synchronization between auditory and motor areas, and asking, crucially, whether this synchronization is modified by different speech rates.

To address this question we computed the PLV between the auditory cortex and speech-motor cortex time series for the different stimulus conditions. To control for artifactual crosstalk between anatomically proximal regions, we calculated the PLV between contralateral areas (Figure 3A upper left panel; see Methods). The PLV for right motor-left auditory and left motor-right auditory signals were calculated and averaged (both PLVs showed the same pattern as the average, see Figure S2A). The synchronization between the auditory and speech-motor brain signals increases for the 2.5, 3.5, and 4.5 syllable/sec conditions, for frequencies around the corresponding stimulus rate. This was not observed for the higher rate conditions (Figure 3A).

**Figure 3.**
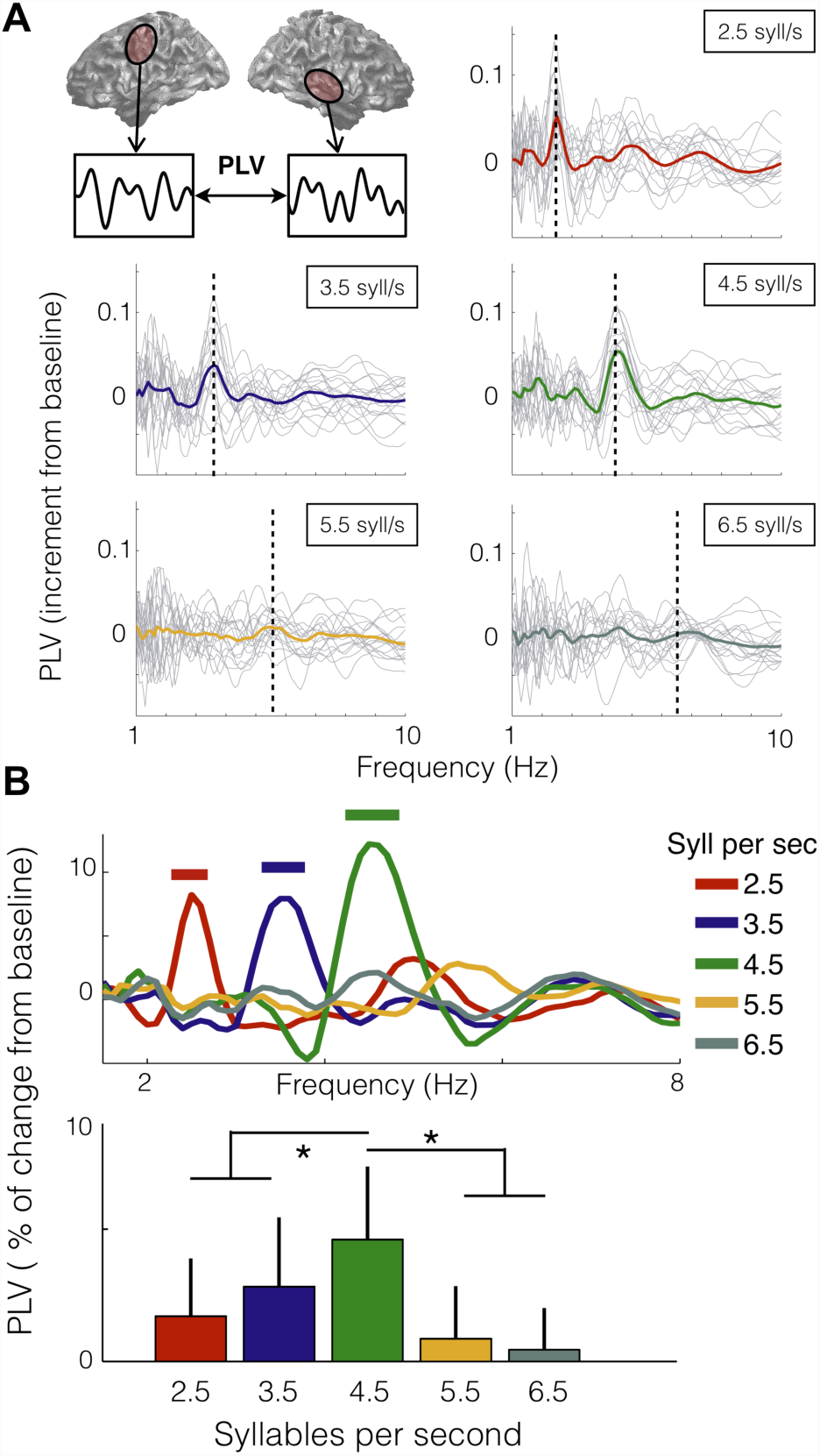
Synchronization between auditory and motor cortices is modulated by the heard syllable rate. A) Increment from baseline of PLV between auditory and motor cortex activity, as a function of frequency. Upper left panel: schematic of analysis. The data panels correspond to the different syllable rate conditions. Light gray lines: individual subject data. Colored line: mean across participants (red: 2.5, blue: 3.5, green: 4.5, yellow: 5.5 and dark gray: 6.5 syllables per seconds). Vertical dashed line: stimulus rate. B) Upper panel: percent change from baseline of PLV as a function of frequency, averaged across subjects. Straight lines on top: significant increment from baseline (Wilcoxon signed rank test p<0.03, FDRcorrected). Lower panel: Mean PLV around the syllable rate of each condition (syllable rate +/- 0.5 Hz). Asterisk: significant difference between conditions (Wilcoxon signed rank test p<0.03, FDR-corrected).

The enhancement of the auditory-motor PLV around the heard syllable rate is statistically significant for the first three conditions (Figure 3B upper panel; Wilcoxon signed rank test p<0.03, FDR corrected). For a higher-resolution characterization of the coupling pattern, we compute the mean PLV, per condition and subject, around the syllable rate (Figure 3B lower panel). There is a significant difference across conditions as shown by a Kruskal-Wallis test (χ^2^(4)=26.5, p<0.001). Unexpectedly, a post-hoc analysis comparing the central condition against all others reveals that the synchronization between areas is enhanced while hearing syllables at 4.5 Hz (Wilcoxon signed rank test p<0.03, FDR-corrected). To test the robustness of this result, we repeated the analysis using a different index to measure coupling (phase lag index, see Methods), and we show that the same synchronization pattern is recovered (Figure S2B). We also performed power analyses for motor and auditory regions; none of these showed an enhancement for the 4.5 syllable rate condition (Figure S3).

To examine whether this observed synchronization pattern is restricted to the interaction between motor and auditory regions, rather than being a generic property of interareal coupling, the PLV pattern between auditory regions across hemispheres was calculated. This auditory-auditory PLV showed peaks at the frequencies corresponding to each syllable rate, but no significant difference across conditions (Figure S4), unlike the auditory-motor PLV pattern.

Next, we tried a different analysis to explore if the strength of the *auditory-motor* coupling depends on the *auditory-speech* coupling. Figure 4 shows the *auditory-motor* PLV as a function of the *auditory-speech* PLV for each syllable rate condition. Fitting a linear regression for each plot reveals that just for the central conditions the correlation between variables is significant (r_2.5_=0.18,p_2.5_=0.33;r_3.5_=0.43, p_3.5_=0.02; r_4.5_=0.49, p_4.5_=0.01;r_5.5_=0,p_5.5_=0.99;r_6.5_=0.3, p_6.5_=0.11). This result suggests that when auditory cortex is synchronized to an external stimulus, this synchronization is conveyed to motor areas only in a restricted frequency range.

**Figure 4.**
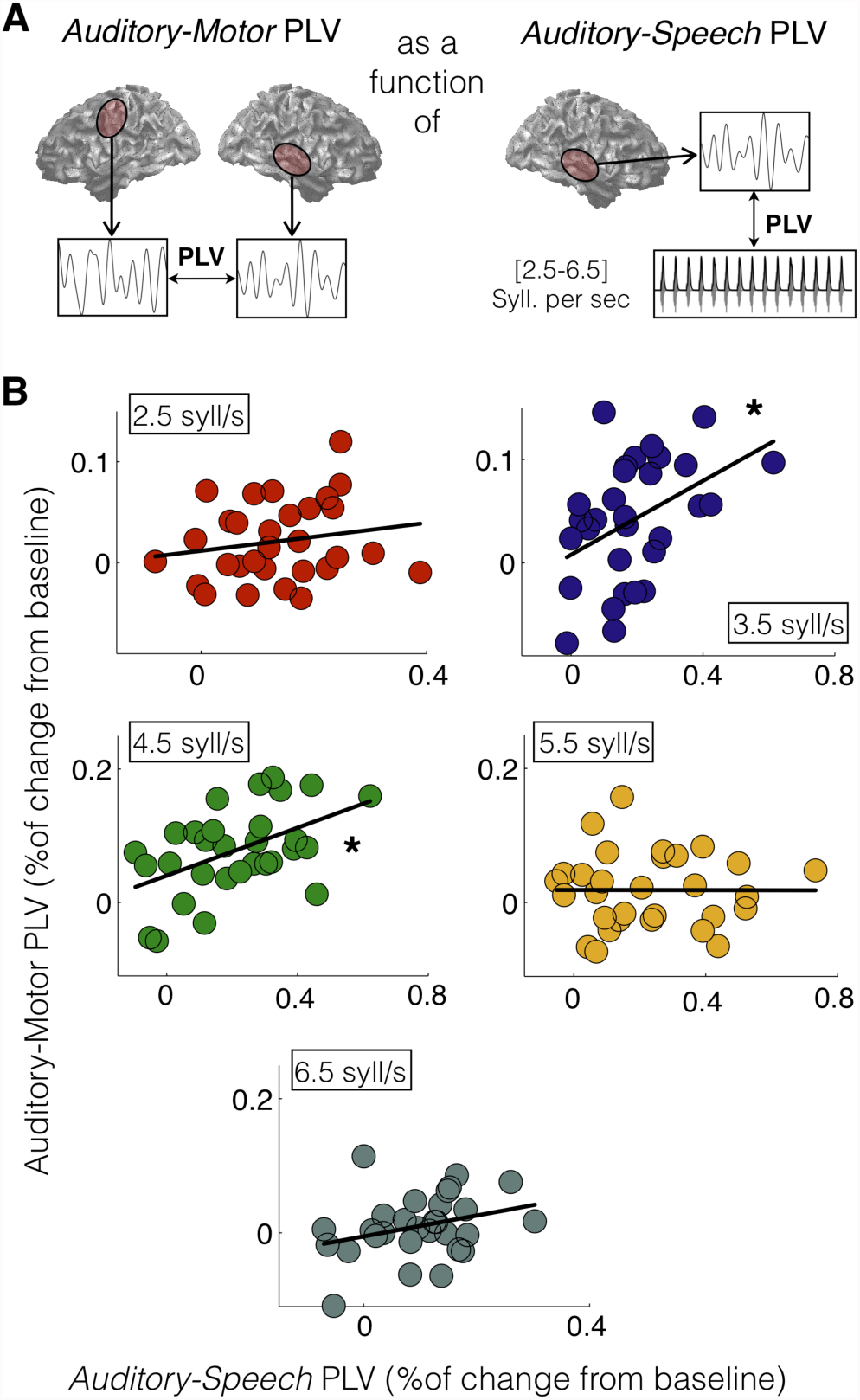
Correlation between auditory-motor PLV and auditory-speech PLV. A) Schematic of analysis.B) Contralateral motor-auditory PLV as a function of the corresponding auditory-stimulus PLV (audL-MotR vs audL and audR-MotL vs audR). Each panel represents a different syllable rate condition (red: 2.5, blue: 3.5, green: 4.5, yellow: 5.5 and dark gray: 6.5 syllables per seconds) and the black line the linear fitting of the data. Asterisk: significant correlation between variables (p<0.05, FDR-corrected).

### A simple neural model successfully accounts for auditory-motor synchronization

Finally, we investigated whether the relatively complex and unexpected auditory-motor synchronization pattern could emerge as a consequence of the underlying neural architecture. We adopted a physiologically inspired neural population model and explored whether it can explain our measurements.

We chose the Wilson-Cowan mean-field approximation to represent speech-motor cortex (Wilson & Cowan 1972). This model is a biophysical model of the interaction between an inhibitory and an excitatory neuronal population that has been widely used in neuroscience (Goldin et al. 2013; Ozeki et al. 2009; Marcos et al. 2013). The model is described by equation (1); where *S* is a sigmoid function whose arguments represent the input activity for each neuron population; *E* and *I* represent the activity of the excitatory and inhibitory neurons, respectively; τ is the membrane time constant; *a* and *b* are synaptic coefficients; *c* and *d* are feedback coefficients, and ϱ is the basal input activity from other areas of the brain.

With respect to the auditory cortex activity (*A*) we assumed it to be entrained by the speech envelope, and we used equation (2) to represent the different experimental conditions. This equation typifies a periodic signal with period *1/syllable rate* that remains at zero for the first 3 seconds.
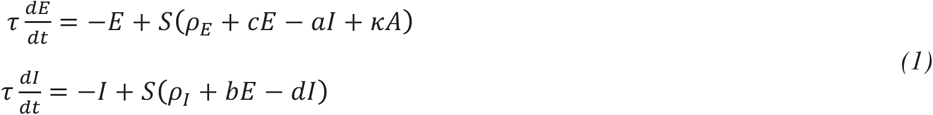

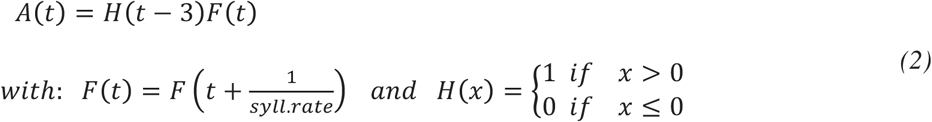

Finally, the interaction between regions takes place because the excitatory motor population receives the auditory cortical activity as input (equation (1) and Figure 5A). The parameter ϰ represent the strength of the interaction.

**Figure 5.**
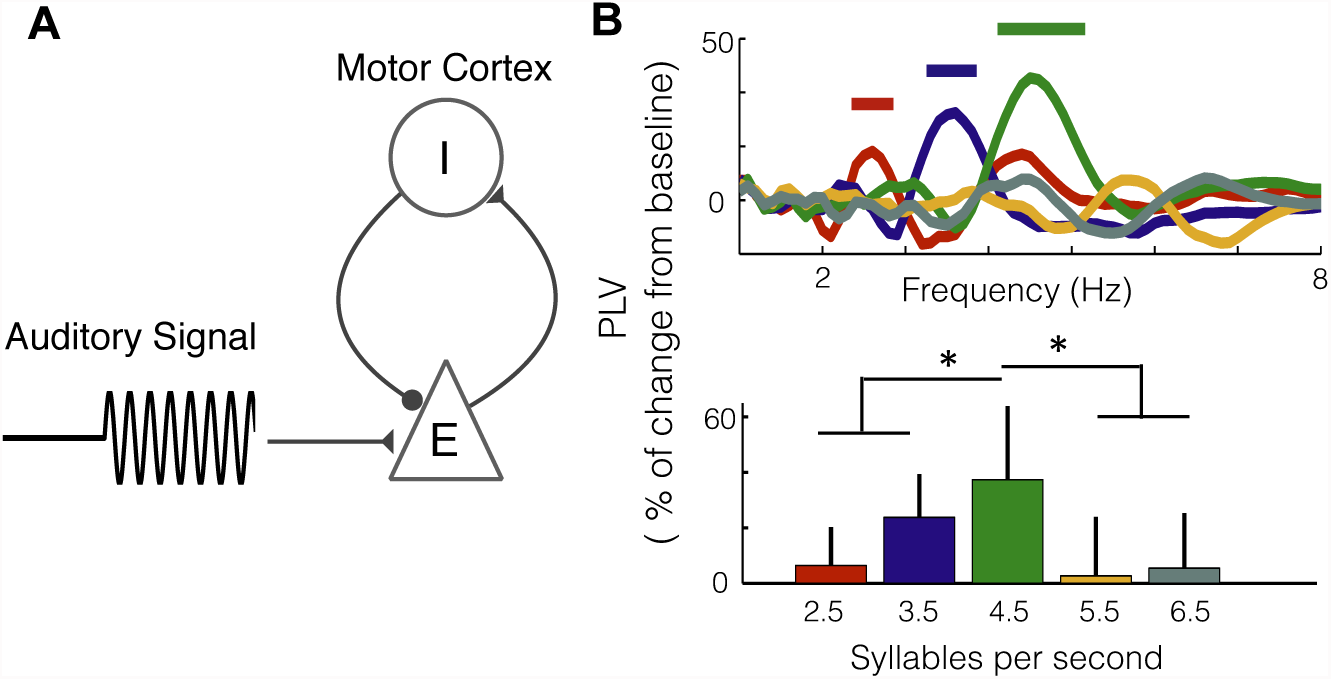
A simple neural model replicates the auditory-motor synchronization pattern. A) Schematic illustration of the neural model: motor cortex modeled through a set of Wilson-Cowan equations representing an inhibitory-excitatory network, the excitatory population receives the auditory cortex activity as input. B) Numerical simulations submitted to the same analyses as the experimental dataset. Straight lines: significant increment from baseline (Wilcoxon signed rank test p<0.03, FDR-corrected). Asterisks: significant differences between conditions (Wilcoxon signed rank test p<0.03, FDR-corrected). Colors identify the different conditions (red: 2.5, blue: 3.5, green: 4.5, yellow: 5.5 and dark gray: 6.5 syllables per seconds).

To reproduce the experimental conditions, the whole MEG dataset for one participant was numerically simulated (see Methods). This methodology allows us to test the neural model, but in doing so also to validate the experimental design and the analyses. With respect to the parameters: *a,b,c,d,*ϱ*_E,_*ϱ*_I_* were fixed within the range typically used in the literature (Hoppensteadt & Izhikevich 1997; Wilson & Cowan 1972); while τ was set to 60 ms in order to get the maximal coupling at 4.5 Hz and ϰ=0.5 to get the desired modulation. It is worth noting that the value of the membrane time constant, τ, is close to the one used in previous works (Marcos et al. 2013; Ozeki et al. 2009) and within the range of experimental results (Tripathy et al. 2015).

The numeric simulations were submitted to the same analyses as the experimental data. The PLVs between the auditory and motor simulated time series, averaged across trials of the same rate, showed the same pattern as the real data (Figure 5B, upper panel). The mean PLV around the condition rate (Figure 5B, lower panel) also displayed the same features as the experimental results. In particular, there was a significant difference across conditions as revealed by a Kruskal-Wallis test (χ^2^(4)=45.6, p<0.001); post-hoc planned comparisons show that the synchronization between regions is enhanced for the 4.5 syllable/sec condition (Wilcoxon signed rank test p<0.03, FDR-corrected).

The numerical simulations show that the experimentally observed synchronization pattern could emerge as a consequence of a relatively simple underlying neural mass model. Moreover, to validate the plausibility of the model, we applied it to a related older study (Black 1951) that addressed auditory speech-motor interaction using delayed auditory feedback. We show that this simple model captures the essential behavioral results of that experiment (Figure S5).

### Behavioral performance does not depend on the syllable rate

We investigated the performance of the individuals on the syllable detection task across the different rate conditions (Figure 6). For all conditions, the performance is above chance (Wilcoxon signed rank test p<0.01, FDR-corrected) and a Kruskal-Wallis test reveals no significant difference across conditions (χ^2^(4)=4.6, p=0.32). This result suggests that the synchronization between areas does not imply a benefit for perception. However, since the present study focuses on exploring the synchronization as a function of the syllable rate, regardless of its functional role, the behavioral task was designed to maintain the individual’s attention on the stimuli - not to explore perception. Further work would be needed to assess the implications of the auditory-motor synchronization on perception.

**Figure 6.**
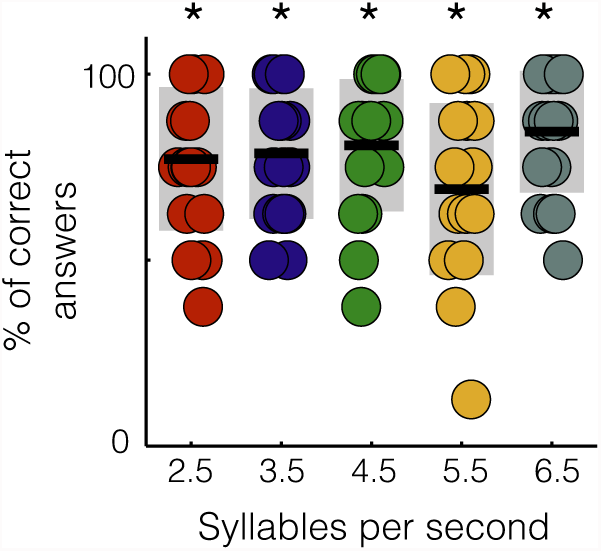
Behavioral results. Performance as a function of the syllable rate condition. Dots: individual participants. Black lines: mean across participants; shadowed region is standard deviation. Asterisks: significant difference from chance (Wilcoxon signed rank test p<0.01, FDR-corrected). Colors identify the different conditions (red: 2.5, blue: 3.5, green: 4.5, yellow: 5.5 and dark gray: 6.5 syllables per seconds).

## DISCUSSION

Our results are best seen in the context of the larger literature on the relationship between perception and action. This area of research, especially the work focused on speech, remains quite controversial (Lotto et al. 2009; Pulvermüller & Fadiga 2010). We address the issue from a new mechanistic perspective. In this neurophysiological experiment, we assess the coupling between auditory and speech -motor regions by testing their synchronization. We exploit the well-documented phenomenon that there is a reliable coupling between acoustic stimuli and auditory cortical activity in order to investigate how sensorimotor synchronization might be implemented across rates. The neural recordings reveal a surprising new phenomenon: whereas the results broadly support the view that there exists a systematic relation between auditory and speech-motor regions, this relationship is not uniform across rates but selectively modulated. Moreover, a simple and well-established neural mass model replicates the experimentally observed auditory-motor synchronization pattern. This invites the interpretation that the auditory-to-speech-motor relation is restricted in phase space as a consequence of the architecture of the underlying neural system.

Typical experimental approaches to study the sensorimotor integration of speech, specifically testing the extent to which motor representations and computations support perception, have pursued multiple lines of evidence, as outlined above. Cumulatively the data on the interaction of auditory and speech-motor cortex suggest some form of interaction but provide no broadly supported functional role - and the mechanisms remain elusive.

We capitalize on a new approach to study the sensorimotor integration of speech. A body of findings from systems and cognitive neuroscience suggests that synchronization between brain regions may represent a crucial contribution to the cortical computational code (Singer 2011). Insofar as anatomically distinct regions can be said to be coupled when measuring their activity, this coupling can be argued to underlie how computations are coordinated. Data from studies on vision, for example, demonstrate how such an approach can inform our hypotheses about the activity underlying higher-order interactions (Bastos et al. 2015). We build on this line of research. The hypothesis that the sensorimotor infrastructure underlying speech may be subserved by synchronized oscillatory activity has not been pursued systematically. Influential data by Park et al. (Park et al. 2015) show the promise of this method when looking more broadly at how synchronization correlates with intelligibility. In light of the promise of this line of attack, our study addresses a fundamental gap in our understanding of synchronization in speech-motor processing. We describe how synchronization comes into play by systematically testing the crucial role of speech rate, and specifically how inter-areal synchronization is modulated by syllable rate. Performance in speech tasks is extremely rate-sensitive, and outside of rates ranging from ~2-9 syllables per second, the ability to extract usable information declines sharply (Ghitza 2014; Dupoux & Green 1997).

Our data show a privileged frequency range for auditory-motor synchronization during perception with a significant enhancement around 4.5 Hz. This enhancement likely reflects a feature of the motor cortex, since the entrainment of auditory areas to stimuli remains stable and more uniform across frequencies. Furthermore, the modeling, too, reveals that adjusting parameters of motor cortex shows the increased synchronization for rates around 4.5 Hz in the simulations. Our findings thus reinforce previous results (Giraud et al. 2007) that reveal frequencies around 4.5 Hz as the natural rhythms of speech motor cortices. This line of reasoning converges, as well, with the fact that it has been shown that mouth movements while speaking occur in the same range of frequencies as highlighted here; notably, the coherence between the speech envelope and mouth movement is also enhanced around 4 Hz (Chandrasekaran et al. 2009). In addition, the mean envelope modulation rate across languages happens to be in the same range (Ding et al. 2017). Altogether, the evidence suggests that the temporal patterns of speech emerge as a consequence of the interaction between cortical areas, with their own intrinsic rhythms, and the additional constraints imposed by the biomechanics of the vocal system consistent with the frame/content theory (MacNeilage 1998).

We systematically investigated the synchronization between auditory and speech-motor cortices across rates and showed that these areas synchronized for a restricted range of frequencies, with a peak at 4.5 Hz. A well-described mean-field neural model of oscillatory neural behavior reproduced our results. To our knowledge, this work is among the first to pursue a mechanistic explanation of how the sensorimotor integration of speech is achieved, regardless of a functional interpretation. The data show that coupling exists between auditory and speech-motor cortex during listening to speech, but that this coupling is rather more restricted than one might have anticipated. It follows that the interaction between auditory and speech-motor regions, in phase space, should be interpreted rather conservatively. This new approach may lead to a more comprehensive understanding of the link between perceptual and motor systems in speech processing and, perhaps, other sensorimotor domains.

## METHODS

### Subjects

Seventeen subjects participated in this study (9 males; mean age 28, range 20-40; 15 native speakers of American English and 2 native speakers of Spanish). All self-reported normal hearing and no neurological deficits, and all had normal structural MRI scans. Participants were paid for taking part in the study and provided written informed consent. Two additional participants completed the experiment but were removed: one because the participant was not able to perform the task and the other because the MEG signal was too noisy. The protocol was approved by the local Institutional Review Board (New York University's Committee on Activities Involving Human Subjects).

### Stimuli

The stimuli consisted of the spoken syllables */ba/*, */wa/*, */ma/* and */va/*; which were synthesized with free online text-to-speech software: *http://www.fromtexttospeech.com/* using a male voice. Each of the syllables was compressed to 120 ms duration using Praat Software (Boersma 2001) and set to the same overall energy (RMS).

Each audio trial of the main experiment consisted of 3 seconds of silence (baseline phase) followed by 6 seconds of syllables (stimulation phase). For each trial, two syllables, the *stable* and the *odd* syllable, were randomly selected from the pool of four (*/ba/*, */wa/*, */ma/* and */va*), to be repeatedly played during the stimulation phase. The syllables were sequentially presented with an occurrence frequency of 0.7 for the *stable* and 0.3 for the *odd*. A silent gap was placed between all syllables in each trial. The inter-syllable silence was uniform within trial, and six different conditions of trials were generated by varying this value between 280 ms and 34 ms. This corresponds to rates of 2.5, 3.5, 4.5, 5.5, 6.5 syllables per second. Eight trials consisting of different syllables were generated for every *syllable rate* condition.

### Tasks

The participants performed the following tasks during MEG recording.

1. *Motor localizer*. Different consonants were sequentially presented on a screen, separated by a stop cue. Subjects were instructed to continuously *mouth* (mimic without vocalizing) each consonant plus an */a/* - vowel as soon as the given consonant was presented on the screen and to continue until the word *stop* was presented. The time between the consonant presentation and the stop cue was jittered between 1.5 and 3 seconds to avoid prediction, and the inter-trial interval (ITI) was jittered between 1 and 1.6 seconds. The consonants used were: */b/*, */v/*, */w/* and */m/*. Twenty-five repetitions of each consonant were presented in a pseudorandomized order.
2. *Auditory localizer*. Single syllables were sequentially played to participants, while they were instructed to stay focused without performing any task. The presented syllables consisted of 25 repetitions of each of the same syllables used to generate the stimuli for the main experiment (*/ba/, /wa/, /ma/* and */va/*). The one hundred total presented syllables were played in pseudo random order, with an ITI jittered between 0.9 and 1.5 seconds.
3. *Main experiment.* During each block, 40 auditory trials (8 trials per syllable rate condition, see *Stimuli*) were played. At the end of each trial one of the four possible syllables (*/ba/, /wa/, /ma/* and */va/*) was displayed on the screen. Participants were instructed to indicate whether the syllable was presented in the trial. The decision was made by pressing a button with the right hand (index finger: *yes, syllable was present;* middle finger: no, *syllable was not present*). The displayed syllable had a 50% probability of being present in the trial. There was no time constraint to make the decision and the next trial started between 0.9 and 1.1 s after button press.

The overall experimental design consisted of 2 blocks of the main experiment followed by the motor localizer, the auditory localizer, and finally 2 more blocks of the main experiment. Two minutes of MEG data were recorded after each participant left the MEG room (*empty room data*). All auditory stimuli were presented binaurally at a mean 75 dB sound pressure, via MEG-compatible tubephones (E-A-RTONE 3A 50Ω, Etymotic Research) attached to E-A-RLINK foam plugs inserted into the ear canal.

### Data acquisition and processing

Neuromagnetic responses were recorded with a 1000 Hz sampling rate using a 157-channel whole-head axial gradiometer system (KIT, Kanazawa Institute of Technology, Japan) in a magnetically shielded room. Five electromagnetic coils were attached to the subject’s head to monitor head position during MEG recording. The coils were localized to the MEG sensors, at three different time points: at the beginning of the experiment, before the motor localizer and before the last two blocks of the main experiment. The position of the coils with respect to three anatomical landmarks: the nasion, and left and right tragus were determined using 3D digitizer software (Source Signal Imaging, Inc.) and digitizing hardware (Polhemus, Inc.). This measurement allowed a coregistration of subjects’ anatomical magnetic resonance image (MRI) with the MEG data. An online bandpass filter between 1 and 200 Hz and a notch filter at 60 Hz were applied to the MEG recordings.

Data processing and analyses were conducted used custom MATLAB code and the FieldTrip toolbox (Oostenveld et al. 2011). For each participant’s dataset, noisy channels were visually rejected. Two procedures were applied to the continuous MEG recordings. First, a least squares projection was fitted to the data from the 2 minutes of *empty room* recorded at the end of each session. The corresponding component was removed from the recordings (Adachi et al. 2001). Second, the environmental magnetic field, measured with three reference sensors located away from the participant's head, was regressed out from the MEG signals using time-shifted PCA (de Cheveigné & Simon 2007). The MEG signals were then detrended and artifacts related to eyeblinks and heartbeats were removed using independent component analysis.

### Structural MRI

High-resolution T1-weighted 3D volume MR data were acquired using a Siemens Allegra 3-Tesla head-only scanner. Each participant’s MRI data were preprocessed following the FieldTrip pipeline. Cortical reconstruction and volumetric segmentation were performed with the FreeSurfer™ image analysis suite.

### Source reconstruction

In order to functionally localize each participant’s auditory and motor areas and to extract the signal coming from those regions it is necessary to reconstruct the cortical current sources generating the magnetic fields recorded by the MEG sensors. Cortically constrained minimum norm estimation (Dale et al. 2000) (MNE) was used. This method approximates the cortical surface as a large number of current dipoles, and estimates the dipole’s amplitude configuration with minimum overall energy that generates the measured magnetic field. Mathematically, it solves the following equation:

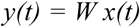

were *x(t)* and *y(t)* are the MEG measurement and the dipole’s amplitudes at time *t*, respectively. *W* is the linear inverse operator defined as:

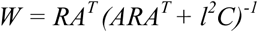

where *R* and *C* represent the spatial covariance matrix of the sources and the noise, respectively, *A* is the linear forward matrix operator that gives the magnetic field at the sensor’s positions produced by orthogonal current unit dipoles, and *l* is a noise scaling factor. The parameters used for the source reconstructions were: the average of the covariance of each trial’s baseline as the noise covariance matrix, a value of 3 for *l* and a source space of 8196 points with a volume conduction model (both reconstructed from each subject’s structural MRI) to compute *A*.

Every current dipole source is a three dimensional vector. When the average across trials is projected back to source-space, power is computed as the norm over the three orientations for each location and time point. If single trials are projected, one signal per source is reconstructed by projecting the three dimensional time-series along the direction explaining the most variance.

#### Auditory area localization

One hundred trials were extracted from the continuous MEG data from the auditory localizer. Trials were defined as 1 second of data centered on the syllable’s onsets. Trials were visually inspected and removed if gross artifacts were detected. Baseline was defined as the 500 ms before syllable onset. Trials were averaged and projected to source-space using MNE. Power was averaged for baseline and the stimulus window (from 50 to 300 ms after the sound onset). Bilateral regions within temporal lobes with activity above baseline (p<0.05 Wilcoxon Rank Test) were identified. Then, for each region, 100 sources around the more active one were selected and defined as the auditory cortex of the corresponding hemisphere.

#### Motor area localization

It has been shown that while speaking, a suppression of the motor cortical 20-Hz rhythm takes place in mouth-related areas (Salmelin et al. 2000). Building on this result, *speech motor areas* were identified by locating this suppression.

Continuous MEG data from the motor localizer protocol was segmented into trials with a duration of 3 seconds (1 second before the starting cue appears on the screen (baseline) and 2 seconds after the mouth-movement period. Each trial was projected back to source-space using MNE, the source signals were filtered between 17 and 25 Hz (μ*-band*), and their amplitudes were computed as the absolute value of the Hilbert transform. Amplitude was averaged for the baseline and stimulus windows (from 500 to 1500 ms after the starting cue). Bilateral regions showing a *μ-band* suppression compared with baseline (p<0.05 Wilcoxon Rank Test) were identified. For each region, 100 sources around the most suppressed one were selected and defined as the *speech motor* cortex of the corresponding hemisphere. Three subjects showed the suppression only in the left hemisphere; therefore no right speech motor area was defined for them.

### Data analysis

#### Main experiment data

As described above, four regions of interest (ROIs) were functionally identified for each participant: right motor (RM), left motor (LM), right auditory (RA) and left auditory (LA). The activity for each ROI while listening to syllables at the different rates was computed in the following analyses.

The data from the four blocks of the Main Experiment protocol were segmented into trials aligned with the auditory presentations. Trials were visually inspected and removed if gross artifacts were detected. Every participant’s data set comprised between 29 and 32 trials for each syllable rate condition (2.5, 3.5, 4.5, 5.5, 6.5 syllables per second). The 3 seconds of silence before the beginning of the syllable repetition were defined as baseline. The cortical activity for each trial was reconstructed using an MNE inverse solution. The signals from the 100 sources comprising each area (LM, RM, LA, RA) were averaged and resampled at 200 Hz, providing one time series per ROI and trial.

The spectrograms of the auditory stimuli were computed using the NSL Auditory Model Matlab Toolbox (Chi T. n.d.) and the stimulus *cochlear* envelopes were calculated by adding the auditory channels between 180 and 7246 Hz. The envelopes were resampled at 200 Hz.

#### Phase locking value

The synchronization between two signals was measured by the phase locking value (PLV) between them as a frequency function. A continuous Morlet wavelet transform (frequency band 1-10 Hz with 0.1 Hz resolution) was applied to both signals the phase evolution for each frequency was extracted and the PLV (as a function of frequency) was computed using the following formula:

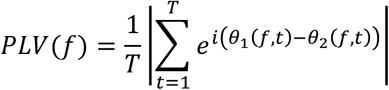

where *f* is the frequency, *t* is the discretized time, *T* is the total number of time points and *θ_1_* and *θ_2_* the phase of the first and the second signals, respectively.

#### Stimulus envelope - brain signal coupling

The PLV was calculated during the stimulation phase of each trial, between each ROI signal and the corresponding auditory stimulus envelope. Windows of 2 seconds length and 1 second overlap were used and the last window was disregarded to avoid boundary artifacts. The results for all time windows were averaged within a trial, providing one PLV per trial. The PLVs were then averaged within each condition and functional ROI (averaging left and right), obtaining one PLV as a function of the frequency for each participant’s data, functional region and for each syllable rate. For each condition the mean auditory PLV around the syllable rate (+/- 0.5 Hz) was computed and compared with the same value evaluated for the baseline data (PLV between baseline auditory time series and the stimulation phase envelope). The percent of change from baseline was estimated as the difference between the PLV during stimulation and during baseline, divided by baseline.

#### Auditory - motor coupling

The PLV for right motor (RM) - left auditory (LA) and left motor (LM) - right auditory (RA) signals were calculated and then averaged. The choice to calculate the PLV between contralateral ROIs was made to avoid artifactual crosstalk between proximal regions(Ghuman et al. 2011). The stimulation phase PLV was computed in the same way as for the stimulus-brain coupling. To estimate the PLV at baseline, one window of 2 seconds right before syllable onset was used and averaged across all trials of a given syllable rate condition.

#### Weighted phase lag index

First, the cross-spectrum between signals was calculated as: 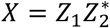, where *Z* represents the Morlet wavelet transform. Then, the debiased wPLI square estimator was computed following Vinck et. al (Vinck et al. 2011), using the following formula:

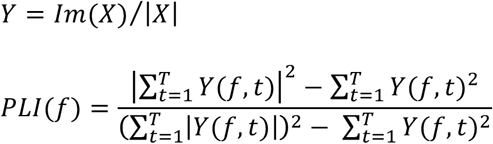

where *f* is the frequency, *t* is the discretized time and *T* is the total number of time points.

#### Model and simulations

The motor cortex activity was described by a minimal model for the dynamics of an inhibitory and an excitatory synaptically-coupled neural populations (Wilson & Cowan 1972). The interaction with auditory areas was modeled by including the auditory cortex activity as an input for the excitatory population as shown in equation (1). The parameters of equation (1) were set as *a*=*b*=*c*=*10* and *d*=*2* according to the literature (Hoppensteadt & Izhikevich 1997). Basal input values were fixed at ρ*_E_*=-1.5 and ρ*_I_*=-3.2; this choice set the system close to an Andronov-Hopf bifurcation, i.e. increasing ρ*_E_* would shift the system from steady state to oscillatory behavior. Finally, ϰ=0.5 and τ=60 ms; to reproduce the experimental synchronization features (an enhancement of the PLV between motor and auditory regions when the last one oscillates at 4.5 Hz).

Auditory cortex activity was modeled by:

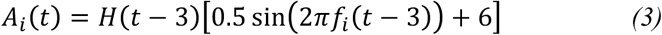

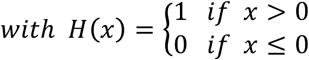

where *t* ∊ [0 9] is time in seconds, *f* is the oscillation frequency in Hz, *i*=1,…,6 represents different condition numbers. The signal emulates the auditory cortex activity while listening to the auditory stimuli: it starts at zero (baseline/silence period) and at *t*=3 (the syllable onset) it increases the basal level and starts oscillating at the syllable rate. The auditory oscillation frequencies reproduce the experimental conditions *f_i_*=[2.5; 3.5; 4.5; 5.5; 6.5] Hz.
For a given auditory activity (*A*, equation 3) the corresponding motor cortex activity (*E-I*) was calculated by numerically solving the set of differential equations (1).

The MEG data simulations were done using the Fieldtrip Toolbox. MEG simulated data for the four blocks of the Main Experiment (32 trials per syllable rate condition) were generated for one subject according to the model. Each ROI activity was simulated by one dipole located in the center of the region. The dipole orientations were random unity vectors and a different set was generated for the last two blocks (half of the data). According to the model, the time courses of the auditory dipoles’ activities were given by *A* and the corresponding (*E-I)* was used for the motor dipoles. The MEG sensors signals were computed using the forward solution derived from the subject's structural image.

For the selected subject, four more blocks of the Main Experiment were included in the experimental protocol, but the audio trials were replaced with silence. The recorded MEG signals were added to the simulated ones as a *background activity*. The mean amplitude of the simulated data during the stimulation phase was set as 2.5% of the amplitude of the *background activity*. The simulated data for the four blocks was submitted to the same processing and analysis as the experimental ones.

## Acknowledgments

We thank Greg Hickok, Greg Cogan, Saskia Haegens and Keith Doelling for comments and advice. The work was supported by NIH 2R01DC05660.

